# Exploring the selenium-over-sulfur substrate specificity and kinetics of a bacterial selenocysteine lyase

**DOI:** 10.1101/2020.02.07.939272

**Authors:** Michael A. Johnstone, Samantha J. Nelson, Christine Van Groesbeck, William T. Self

## Abstract

Selenium is a vital micronutrient in many organisms. While traces are required for utilization by the microbe, excess amounts are toxic; thus, selenium can be regarded as a biological “double-edged sword”. Selenium is chemically similar to the essential element sulfur, but curiously, evolution has selected the former over the latter for a subset of oxidoreductases. Enzymes involved in sulfur metabolism are less discriminate in terms of preventing selenium incorporation; however, its specific incorporation into selenoproteins reveals a highly discriminate process that is not completely understood. We have identified SclA, a selenocysteine lyase in the nosocomial pathogen, *Enterococcus faecalis*, and characterized its enzymatic activity and specificity for L-selenocysteine over L-cysteine. It is known that a single residue in the human selenocysteine lyase, D146, is considered to control selenocysteine specificity. Thus, using computational biology, we identified H100, a D146 ortholog in SclA, and generated mutant enzymes with site-directed mutagenesis. The proteins were overexpressed, purified, and characterized for their biochemical properties. All mutants exhibited varying levels of activity towards L-selenocysteine, suggesting a catalytic role for H100. Additionally, L-cysteine acted as a competitive inhibitor towards all enzymes with higher affinity than L-selenocysteine. Our findings offer key insight into the molecular mechanisms behind selenium-over-sulfur specificity and may further elucidate the role of selenocysteine lyases *in vivo*.

---

Selenium is an important trace element found in all known domains of life. In many organisms, selenium functions as an essential micronutrient when present in small amounts, but in excess, the reactive nonmetal becomes highly toxic and therefore must be kept under tight regulation—this phenomenon is why selenium is often regarded as a “double-edged sword” in biology. Selenium shares many properties with the nonmetal, sulfur, as both elements are part of the chalcogens group on the periodic table. While both elements are similar in terms of atomic character and involvement in redox reactions, sulfur is much more abundant in nature as a macronutrient.

The most well-characterized biological form of selenium is selenocysteine, the 21^st^ amino acid (1,2). Unlike the standard twenty proteinogenic amino acids, selenocysteine is genetically encoded by the UGA stop codon, which is reprogrammed for selenocysteine insertion into the nascent polypeptide by tRNA^Sec^ and a stem loop structure termed the SECIS element (<u>se</u>leno<u>c</u>ysteine insertion <u>s</u>equence) (3,4). In bacteria, this process of specific incorporation is mediated by the following gene products: selenocysteine synthase (SelA), selenocysteine-specific elongation factor (SelB), tRNA^Sec^ (*selC*), and selenophosphate synthetase (SelD) (2,5,6). Interestingly, selenocysteine possesses chemical attributes that are not present in its sulfur analog, cysteine. First, the selenol group of selenocysteine has a p*K*_a_ of ∼5.2 while the thiol group of cysteine has a p*K*_a_ of ∼8.5; because of this, free selenocysteine is deprotonated at physiological pH rendering it extremely reactive compared to cysteine’s protonated side chain (7). Second, selenocysteine can resist over-oxidation in some redox-dependent enzymes such as thioredoxin reductase while cysteine usually cannot recover from such a destructive event (8). Lastly, the presence of a selenocysteine residue in the active sites of redox-active enzymes generally confers a catalytic advantage compared to their cysteine-containing homologs and mutants (9-11). In addition to selenoproteins, selenium can also be incorporated into the wobble position of some tRNAs as 2-selenouridine, but the role of this modification is poorly understood (12-14). Finally, selenium can also be incorporated as a labile cofactor into the active sites of certain molybdenum hydroxylases, often playing a role in enzyme catalysis (15-18). These subtle properties regarding selenium can be seen as evolutionary favorable in the context of some biological systems.

While the aforementioned redox advantages of selenium imply a vast difference between it and sulfur, biology often has a difficult time distinguishing between the two atoms. When cells contain high selenium-to-sulfur ratios, selenium is often misincorporated into biomolecules in place of sulfur by key sulfur metabolic enzymes which are non-discriminate and fail to prevent the insertion of selenium into these molecules—this mechanism is still not completely understood (19). In bacteria, these key enzymes are *O*-acetylserine sulfhydrylase (CysK) and cysteinyl-tRNA synthetase (CysRS). CysK respectively generates cysteine and selenocysteine using sulfide (HS^-^) and selenide (HSe^-^), while CysRS can mischarge tRNA^Cys^ with selenocysteine instead of cysteine (19,20). Therefore, via the non-specific incorporation pathway, the cysteines and methionines of proteins are inadvertently replaced by selenocysteines and selenomethionines, respectively (19,21-25). These misincorporation events could likely play a role in selenium toxicity. Nevertheless, many organisms require selenium for life, so it seems logical that the specific incorporation pathway functions to provide the trace element to fill that critical need in the presence of immense amounts of sulfur.

While the mechanisms for acquiring and positioning selenium for SelD-dependent activation to selenophosphate (SePO_3_) are poorly characterized, a class of proteins has been identified as a likely candidate for this purpose. NifS-like proteins are pyridoxal 5’-phosphate (PLP)-dependent aminotransferases, homologous to the *nifS* (<u>ni</u>trogen fixation) gene product in *Azotobacter vinelandii*. (26). This enzyme functions as a cysteine desulfurase (CDS) where it degrades L-cysteine into L-alanine and HS^-^ (26,27). After L-cysteine degradation, the sulfur atom is covalently bound to the catalytic thiolate of the CDS through a persulfide intermediate, and with the help of an array of scaffolding proteins, the inorganic sulfur is subsequently mobilized to important biomolecules such as iron-sulfur (Fe-S) cluster-containing proteins, thiamine, and molybdenum cofactor, demonstrating a clear role for CDSs *in vivo* (27). The majority of knowledge stems from analyses on the known CDSs in *Escherichia coli*: IscS, CSD (CsdA), and SufS (CsdB). While the three enzymes recognize L-cysteine as their primary substrate, they also nonspecifically act on L-selenocysteine with varying degrees of activity (27-31).

On the other hand, a subset of NifS-like enzymes acts specifically on L-selenocysteine with no regard to the L-cysteine pool within the cell, thus exhibiting selenocysteine lyase (SCL) activity and therefore degrading L-selenocysteine into L-alanine and HSe^-^ (32,33). Moreover, SCLs are curiously inhibited by L-cysteine in either a competitive or non-competitive manner, depending on the enzyme (32-34). SePO_3_, the selenium donor for selenoprotein biosynthesis, is generated from HSe^-^ via SelD (35,36). Because SCLs generate HSe^-^, it is possible that they may be involved in channeling selenium into this pathway, analogous to sulfur mobilization by CDSs. This interaction has been demonstrated *in vitro* with mammalian SCL (37,38). Selenium channeling by this enzyme may be primarily mediated by its selenium-over-sulfur discriminatory ability. Unfortunately, this is speculative without clear evidence of the exact mechanism by which this occurs. While the mammalian enzyme has been reported to be involved in glucose, lipid, and amino acid metabolism in mice (39-41), the *in vivo* role of this subclass of NifS-like enzymes, especially in prokaryotes, is still unclear. The lack of information stems from the fact that most of the SCLs that have been biochemically characterized are from mammals where selenium metabolism is not as well-understood as it is in prokaryotes (32,34,42). In fact, SCL activity has been reported in several prokaryotes, but only one of them was purified and characterized from the crude lysate of *Citrobacter freundii* (33,43). Indeed, characterizing the genetic products of open reading frames that encode putative SCLs would help reveal more about the biological role of these enzymes.

While studying the selenium metabolism of several pathogens, our group identified a NifS-like protein in *Enterococcus faecalis* (44). *E. faecalis* is a multidrug-resistant pathogen and a prominent nosocomial responsible for endocarditis, bacteremia, and urinary tract infections (45). Aside from these diseases, its alarming ability to effortlessly form biofilms on the surfaces of medical devices such as catheters and pacemakers makes it an ongoing healthcare issue (46). Curiously, *E. faecalis* contains an “orphan *selD*” where it does not contain the genes necessary to synthesize selenoproteins or selenium-containing tRNAs but instead may utilize SePO_3_ for some other unknown trait (44,47). Additionally, genes that are putatively involved with the biosynthesis of a selenium-dependent molybdenum hydroxylase (SDMH) were identified in the same genetic locus as *selD* (44,48).

We have previously demonstrated that the SDMH in question is a xanthine dehydrogenase (XDH) that has been shown to significantly increase biofilm formation in the presence of uric acid, molybdate, and selenite (49). In the same genetic locus, many uncharacterized genes were annotated to be hypothetically involved in selenium metabolism—one of which is EF2568, whose gene product encodes a putative NifS-like protein (44). Based on our initial characterization of the protein and the fact that it may be involved in selenium trafficking according to its genetic location amongst other selenium-related genes, we hereby propose a renaming of the open reading frame to *sclA* (<u>s</u>eleno<u>c</u>ysteine <u>l</u>yase <u>A</u>). Its true role *in vivo*, however, is still unclear, and the mechanism by which it specifically recognizes selenium is unknown.

An investigation into the selenium-over-sulfur specificity mechanism of human SCL (hSCL) has shed some light on this topic. A site-directed mutagenesis approach found that mutation of the aspartic acid at amino acid position 146 (D146) to a lysine (D146K) resulting in novel activity towards L-cysteine (50,51). This implied that D146 is either the sole determinant of selenium specificity or at least plays a significant role in the mechanism. Specifically, the authors proposed that, while a protonated thiol could not react with the thiol of L-cysteine, the nucleophilic selenolate anion of L-selenocysteine may deprotonate the catalytic cysteine residue’s thiol which could then attack the electrophilic selenol group, causing the reaction to proceed with high specificity for selenium (51). In this case, it is believed that the nearby carboxylate anion of D146 serves to increase the p*K*_a_ of C388 (the catalytic cysteine) through a like-charge destabilization of its thiolate anion; thus, the protonated thiol species would dominate, allowing only reaction with L-selenocysteine. When D146 was mutated to a lysine, the charge was inverted and may have reduced the p*K*_a_ of C388. It is possible that the lysyl side chain’s amino cation stabilized the deprotonated thiolate species, allowing C388 to react with both substrates. Overall, this implies that the presence of D146 heavily influences the p*K*_a_ of C388 and may prevent the residue from achieving the thiolate form present in other NifS-like enzymes. It should be noted, however, that the p*K*_a_ of C388 was not determined in this study.

Because D146K of hSCL was reported to eliminate L-selenocysteine specificity, we wondered if an orthologous residue in SclA shared a similar role in influencing the p*K*_a_ of its catalytic cysteine. In this work, we set out to determine the function of the ortholog H100 through a site-directed mutagenesis approach based on the previous work on hSCL. We hope that the results of this work further add knowledge to the mechanism of selenium-over-sulfur discrimination and give insight as to how SCLs play a role in trafficking selenium and preventing toxicity.

## Results

### H100 of SclA aligns to D146 of hSCL

It is known that the selenocysteine specificity of hSCL is strongly governed by D146 (50,51). With this in mind, we wondered if this “single-residue” property held true in SclA. To search for putative specificity-determining residues in SclA, we generated a multiple sequence alignment with various characterized NifS-like proteins using Clustal Omega (Fig. 1). According to the alignment, H100 appeared to directly align with D146, implying that H100 may likewise function as a determinant for selenium-over-sulfur discrimination. Furthermore, various conserved regions were shared throughout the protein sequences including the catalytic cysteines, PLP-binding lysines, and group I and II bacterial NifS-like consensus sequences. In the case of SclA, the presence of the RXGXHCA consensus sequence in the active site confirms its identity as a group II bacterial NifS-like protein (27,29). These aligned regions further added confidence to the hypothesis that H100 may function similarly to D146.

**Figure 1:**
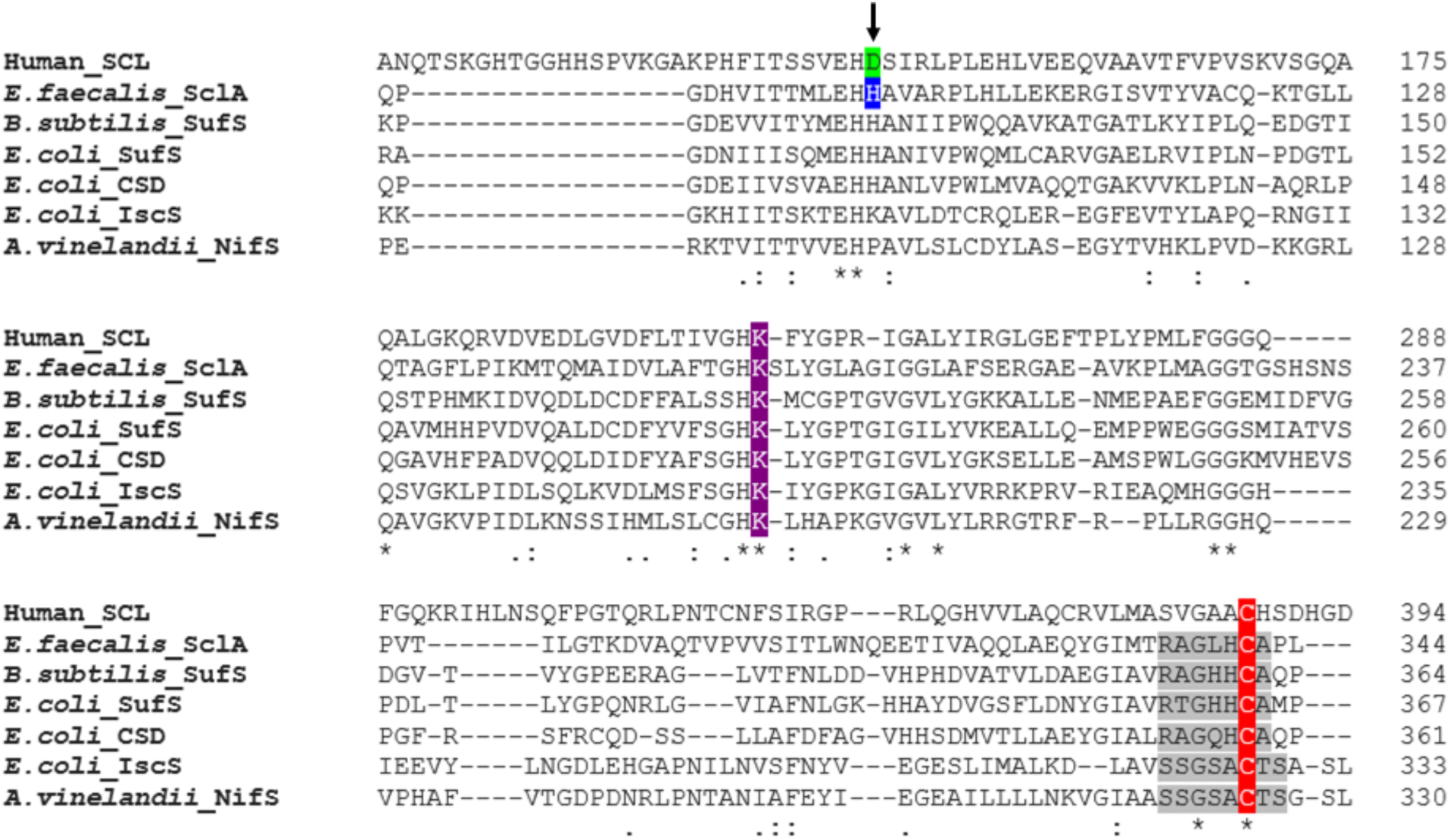
Sequence alignment of SclA and other NifS-like proteins. NifS-like protein sequences from several organisms were aligned using Clustal Omega (70). Smaller segments of the alignment were selected to highlight relevant residues. The arrow indicates the alignment of D146 of human SCL (green) to H100 of *E. faecalis* SclA (blue). For each enzyme, the core catalytic cysteine residue is highlighted in red while the PLP-binding lysine residue is highlighted in purple. The consensus sequences for group I (SSGSACTS) and group II (RXGXHCA) bacterial NifS-like proteins are shaded in gray (27,29). Fully conserved residues are indicated by asterisks (**\***). Partially conserved residues are indicated by colons (:) for residues with strong similarity and periods (.) for residues with weak similarity.

After the initial sequence alignment, we were interested in comparing the structural positions of D146 and H100—while the former is a verified catalytic residue in the active site, the location of the latter is unknown. To further investigate the potential role of H100 as a specificity-determining residue and probe the structural question, we generated three-dimensional models of SclA and hSCL using PyMOL. While hSCL could easily be modeled using its known crystal structure, there was greater difficulty in modeling SclA since it lacked one; thus, we used SWISS-MODEL to search for closely related structural homologs of SclA. The top result was SufS from *Bacillus subtilis* which shared 36.99% sequence identity with the *E. faecalis* enzyme; furthermore, a crystal structure of SufS had been determined in a previous study (52). With SWISS-MODEL, we threaded the primary sequence of SclA through SufS, generated a putative 3-D structure, and further modeled it using PyMOL. The results of both homology models are shown in Fig. 2. For this section, the 4’-deoxypyridoxine phosphate (PLR) ligand present in the hSCL PDB file is referred to as “PLP” for simplicity. The model of wild-type (WT) hSCL highly resembled its crystal structure, containing the conserved NifS-like residues—the PLP-bound K259 and catalytic C388—in its active site (Fig. 2A). Another conserved residue, H145, was located near the PLP molecule and C388, implying a role in catalysis. Finally, D146 was found within 4 angstroms of C388, as predicted by the crystal structure (50). Additionally, Fig. 2B shows the predicted active site of hSCL with a D146K mutation, visualizing the difference in charge. In Fig. 2C, the predicted active site of WT SclA contained the following conserved NifS-like residues: K202 and C341. Because this structure is purely hypothetical, PLP could not be modeled bound to K202 like it was bound to K259 of hSCL, having already been present in the latter’s crystal structure. The residue that aligns to H145 from hSCL in SclA, H99, appeared to likewise position itself next to C341. Furthermore, H100 also seemed to occupy a position away from the cluster of residues similar to D146 of hSCL. The spatial configuration of these residues in SclA (Fig. 2C) seems to mimic the configuration in hSCL (Fig. 2A). While the bacterial model is strictly putative, these data imply that the active sites of both enzymes are highly comparable.

**Figure 2:**
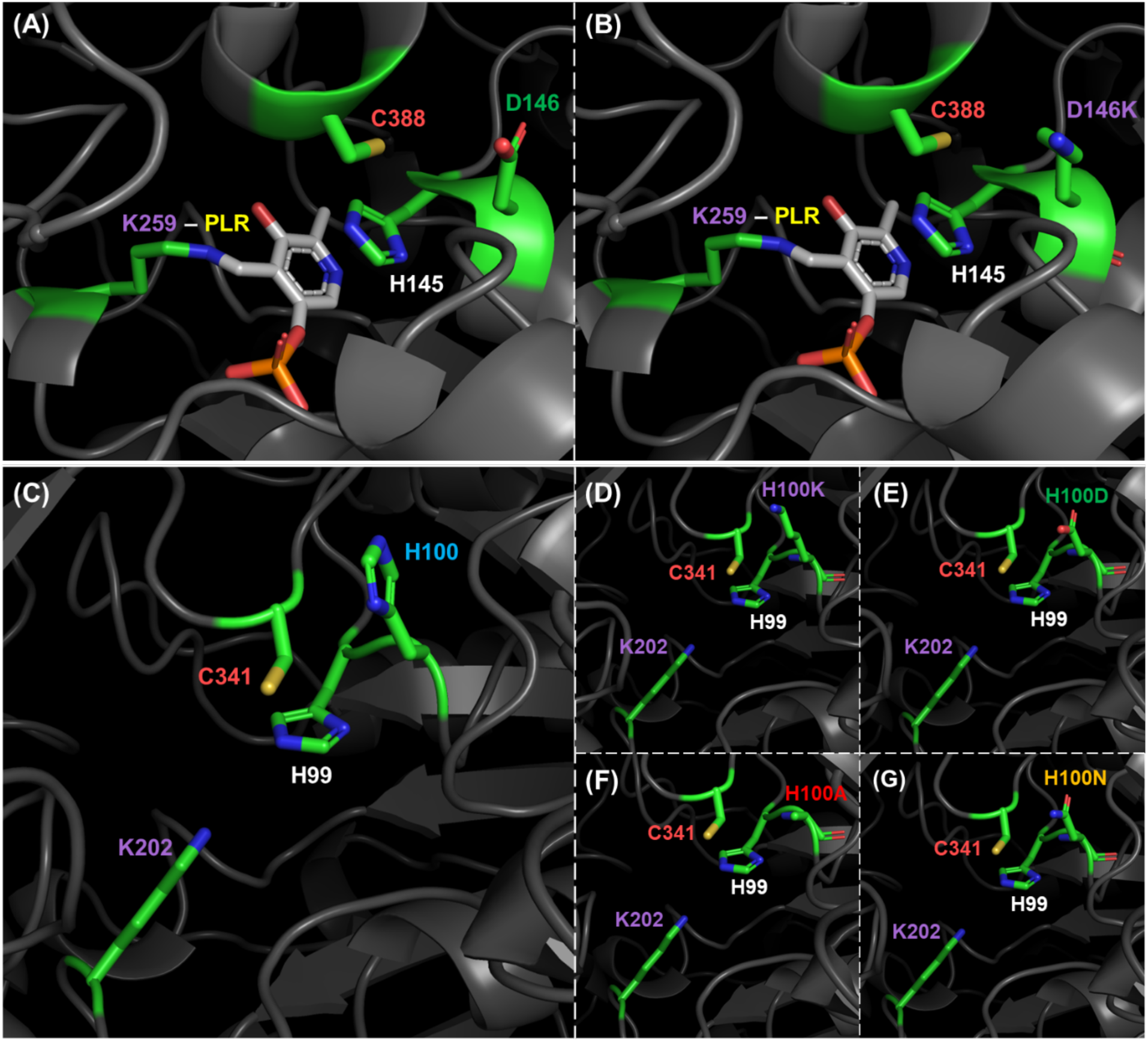
The modeled structure of SclA is similar to the defined hSCL active site. Three-dimensional structures of human SCL and SclA were modeled using PyMOL (72). The primary sequence of SclA was threaded through the established crystal structure of *B. subtilis* SufS as a template to create a homology model using SWISS-MODEL (73). **(A)** Active site of WT human SCL as determined by X-ray crystallography (50). **(B)** Active site of human SCL with a D146K mutation. **(C)** Putative active site of WT SclA, **(D)** SclA H100K, **(E)** SclA H100D, **(F)** SclA H100A, and **(G)** SclA H100N, based on homology modeling with SufS using PyMOL.

### H100 point mutants of SclA were successfully generated

Since a D146K mutation in hSCL resulted in a change in substrate specificity, we hypothesized that a mutation of H100 would influence SclA in a similar manner. To begin probing this question, H100 was mutated to several amino acid residues of differing character (H100K, H100D, H100A, and H100N) via ‘Round-the-horn’ site-directed mutagenesis (53,54); these mutations were selected to ascertain the substrate specificity and catalytic influence of basic, acidic, nonpolar, and polar uncharged properties, respectively. Additionally, the hypothetical active sites of these mutants were modeled similarly using PyMOL (Fig. 2D-2G). An ethidium bromide agarose gel showcasing the individual ∼6.9 kb PCR products after mutagenesis is shown in Fig. S1A, clearly demonstrating successful PCRs and product purity. These PCR products were gel-extracted, ligated, and transformed into *E. coli* strain NEB 5-α which was then used for plasmid minipreps. To confirm the presence of each mutation, the mutant plasmids were sequenced (Fig. S1B). The sequencing chromatograms for WT and all mutants revealed well-defined signal peaks with minimal background noise and clear distinction between the base calls, indicating the purity of the sequencing reactions and successful mutagenesis of H100.

### SclA H100 mutants can be overexpressed and purified

In order to analyze the substrate specificity of SclA, soluble proteins were required for our kinetics assays. All proteins were overexpressed and purified in the same manner as described in Materials and Methods. Thus, Fig. S2A, S2B, and S2C were included to showcase only the H100K purification as an example and avoid redundancy. In Fig. S2A, a relatively sizable ∼42 kDa band corresponding to the theoretical weight of SclA appeared after induction with IPTG, indicating that SclA can be overexpressed heterologously; moreover, successful induction of the target protein further confirmed the results of the earlier transformation. Additionally, the protein seemed to present mostly within the insoluble fraction even after a 7-hour induction at 25 °C with 0.5 mM IPTG. Nevertheless, the amount of target protein in the soluble lysate was sufficient for purification and downstream experiments. Initially, cobalt affinity chromatography was used to capture SclA from the lysate, successfully eliminating most of the *E. coli* contaminants with 5 mM imidazole in the batch-binding step and 20 mM imidazole in subsequent washes (Fig. S2B). Contaminants with molecular weights lower than ∼40 kDa co-eluted with the target protein—the banding pattern implies either proteolytic degradation or N-terminal truncation since all mutants were N-terminally His_6_-tagged. Finally, anion exchange chromatography was used as a polishing step to rid the target of any lingering contaminants (Fig. S2C). After purification was completed, elutions were pooled together, dialyzed, and inspected on an SDS-PAGE gel (Fig. S2D). All results described for purification of this mutant were comparable for all other purifications, and all mutants in Fig. S2D were determined to be pure for downstream experimentation. Overall, these results demonstrate that SclA and selected point mutants can be overexpressed and purified to homogeneity for future studies; additionally, because the H100 mutants could be purified from the soluble lysates, it stands to reason that H100 does not seem to play a significant role in protein folding.

### SclA H100 mutants demonstrate varying Michaelis-Menten kinetics for L-selenocysteine

To ascertain the effects of H100 mutations on substrate specificity, purified proteins were assayed for SCL activity using the lead acetate assay. Because the formation of a persulfide linkage between the sulfur atom and the catalytic cysteine is reported in other NifS-like enzymes, a similar mechanism may be happening in SCLs (55). In SCLs, HSe^-^ may not be the “true” product of the reaction; rather, the selenium atom may be linked to the catalytic cysteine through a selenosulfide bond similar to CDSs. In order to break this bond and release the selenium, a strong reductant like dithiothreitol (DTT) must be used. Moreover, HSe^-^ can only be detected spectrophotometrically if reduced to H_2_Se and complexed with lead, forming PbSe which is observable at 400 nm; thus, the catalytic activity of SclA can be analyzed with the lead acetate assay in order to determine the kinetics of its SCL activity.

A thorough investigation of each enzyme revealed that they all exhibited Michaelis-Menten kinetics for L-selenocysteine at varying levels relative to each other. The kinetic parameters of each mutant are catalogued in Table 1. WT SclA exhibited a *K*_m_^app^ of 3.55 ± 0.32 mM for L-selenocysteine while H100K and H100N displayed similar values of 3.71 ± 0.17 mM and 3.38 ± 0.26 mM, respectively (Fig. 3). Curiously, H100D displayed the weakest affinity for L-selenocysteine, showcasing a *K*_m_^app^ of 5.02 ± 0.49 mM (Fig. 3); conversely, H100A demonstrated the strongest affinity for L-selenocysteine with a *K*_m_^app^ of 1.86 ± 0.13 mM (Fig. 3). Furthermore, H100K and H100N exhibited the highest *V*_max_ ^app^ values of 8.34 ± 0.13 µmol min^-1^ mg^-1^ and 9.89 ± 0.25 µmol min^-1^ mg^-1^ while WT, H100D, and H100A indicated relatively similar lower values. Though H100K and H100N exhibited the highest significant turnover numbers (*k*_cat_) of 5.79 ± 0.09 s^-1^ and 6.86 ± 0.17 s^-1^ respectively, the most catalytically efficient (*k*_cat_/*K*_m_^app^) enzymes were H100A with a value of 2.44 s^-1^ mM^-1^ and H100N with the value of 2.03 s^-1^ mM^-1^ (Table 1). While H100N exhibited faster turnover, the higher binding affinity for L-selenocysteine in H100A heavily contributed to its catalytic efficiency. However, all significant kinetic differences regarding *K*_m_^app^, *V*_max_^app^, *k*_cat_, and *k*_cat_/*K*_m_^app^ according to their H100 mutations only resulted in <2-fold changes compared to WT.

**Table 1:** Kinetic parameters of SclA and selected point mutants.

| Enzyme | $K_m^{\text{app}}$<br>(mM L-Sec) | $V_{\text{max}}^{\text{app}}$<br>($\mu\text{mol min}^{-1} \text{mg}^{-1}$) | $k_{\text{cat}}$<br>($\text{s}^{-1}$) | $k_{\text{cat}}/K_m^{\text{app}}$<br>($\text{s}^{-1} \text{mM}^{-1}$) | $K_i^{\text{app}}$<br>(mM L-Cys) |
| --- | --- | --- | --- | --- | --- |
| **WT** | $3.55 \pm 0.32$ | $6.38 \pm 0.25$ | $4.43 \pm 0.17$ | 1.25 | $0.55 \pm 0.09$ |
| **H100K** | $3.71 \pm 0.17$ | $8.34 \pm 0.13$ | $5.79 \pm 0.09$ | 1.56 | $1.24 \pm 0.15$ |
| **H100D** | $5.02 \pm 0.49$ | $6.50 \pm 0.30$ | $4.52 \pm 0.21$ | 0.90 | $0.62 \pm 0.07$ |
| **H100A** | $1.86 \pm 0.13$ | $6.53 \pm 0.14$ | $4.54 \pm 0.10$ | 2.44 | $0.39 \pm 0.05$ |
| **H100N** | $3.38 \pm 0.26$ | $9.89 \pm 0.25$ | $6.86 \pm 0.17$ | 2.03 | $0.57 \pm 0.06$ |

Standard deviations were calculated by GraphPad Prism 8 and are presented as “ ± SD”. $K_i^{\text{app}}$ constants were determined separately from all other kinetic constants as outlined in Materials and Methods.

**Figure 3:**
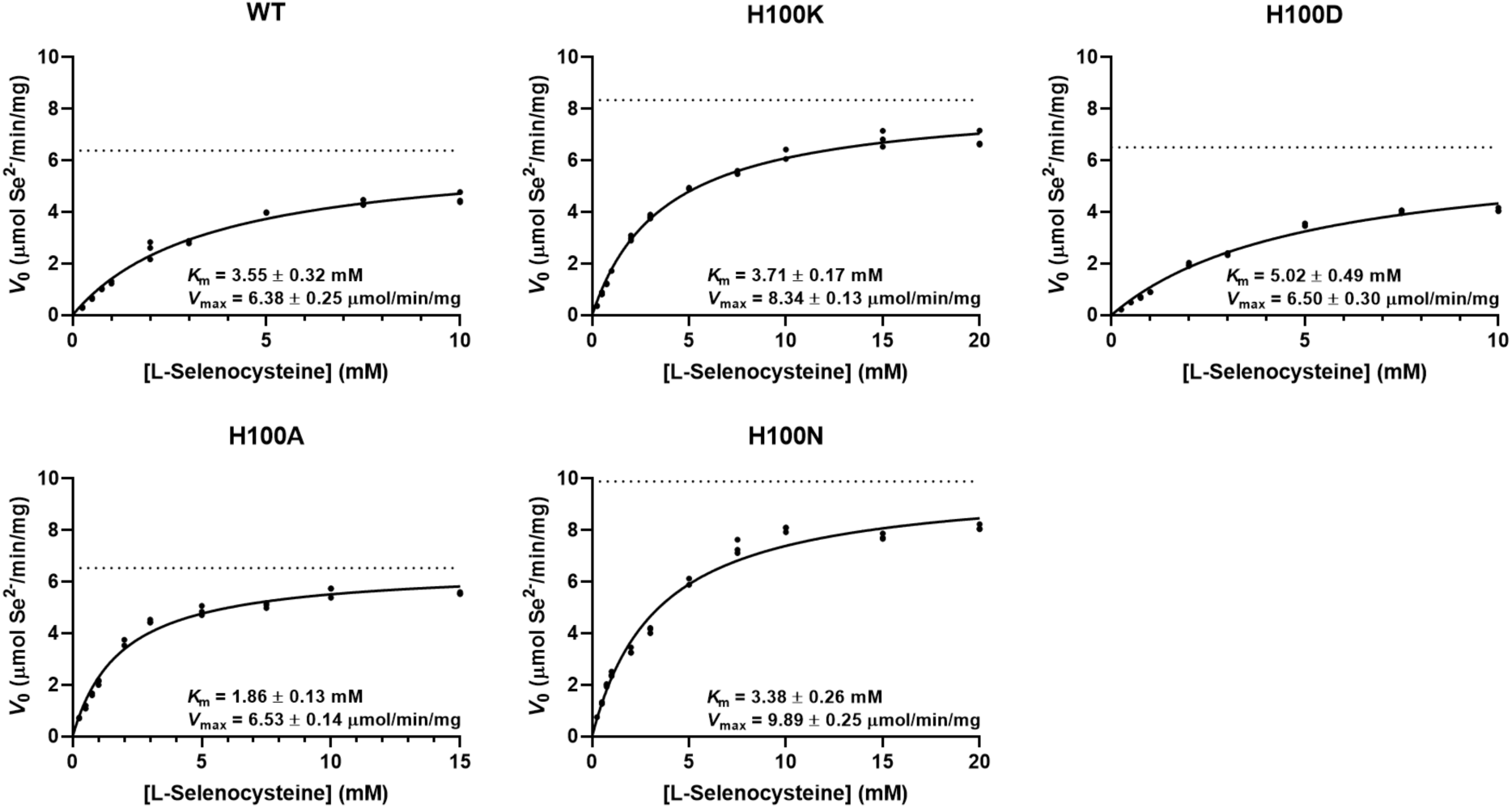
SclA and H100 mutants exhibit varying Michaelis-Menten kinetics with L-selenocysteine as a substrate. The selenocysteine lyase kinetics of purified WT SclA and H100 mutants were determined using the lead acetate assay. The saturation curves were generated by fitting initial rates (*V*_0_) to the Michaelis-Menten equation using GraphPad Prism 8. Kinetic parameters provided below the curves are described further in Table 1. *V*_max_^app^ values are indicated by dotted asymptotes drawn on the y-axes. Data points above 10 mM (for WT and H100D) and 15 mM L-selenocysteine (H100A) were excluded due to apparent substrate inhibition.

It is worth noting that some enzymes displayed substrate inhibition at concentrations above 10 mM L-selenocysteine. The enzymes that experienced substrate inhibition were WT, H100D, and H100A. Because this non-Michaelis-Menten behavior skewed the Michaelis-Menten parameters of the data sets, these inhibitory concentrations were omitted from their respective figures. Altogether, the kinetic variance in each mutant confirms that H100 is in the active site of the enzyme and may be playing a role in catalysis.

### L-Cysteine inhibits the SCL activities of H100 mutants

It is known that L-cysteine acts as an inhibitor for SCLs (32-34). To determine the effect of L-cysteine on the SCL activity of the bacterial enzyme, the lead acetate assay was repeated with several concentrations of L-cysteine for WT and all mutants. Double-reciprocal plots were generated from the initial rates of HSe^-^ formation to determine the type of inhibition mediated by L-cysteine (Fig. 4). Upon visible inspection of the double-reciprocal plot for WT, L-cysteine seemed to act as a competitive inhibitor due to the fact that the trendlines appear to roughly converge on a single point above the x-axis while the slopes change with each varying condition, resulting in different x-intercepts. Thus, the *V*_max_^app^ does not appear to change while the *K*_m_^app^ changes significantly upon L-cysteine titration, indicating competitive inhibition. While this effect differs slightly with each mutant, the overall trend appears to be the same.

**Figure 4:**
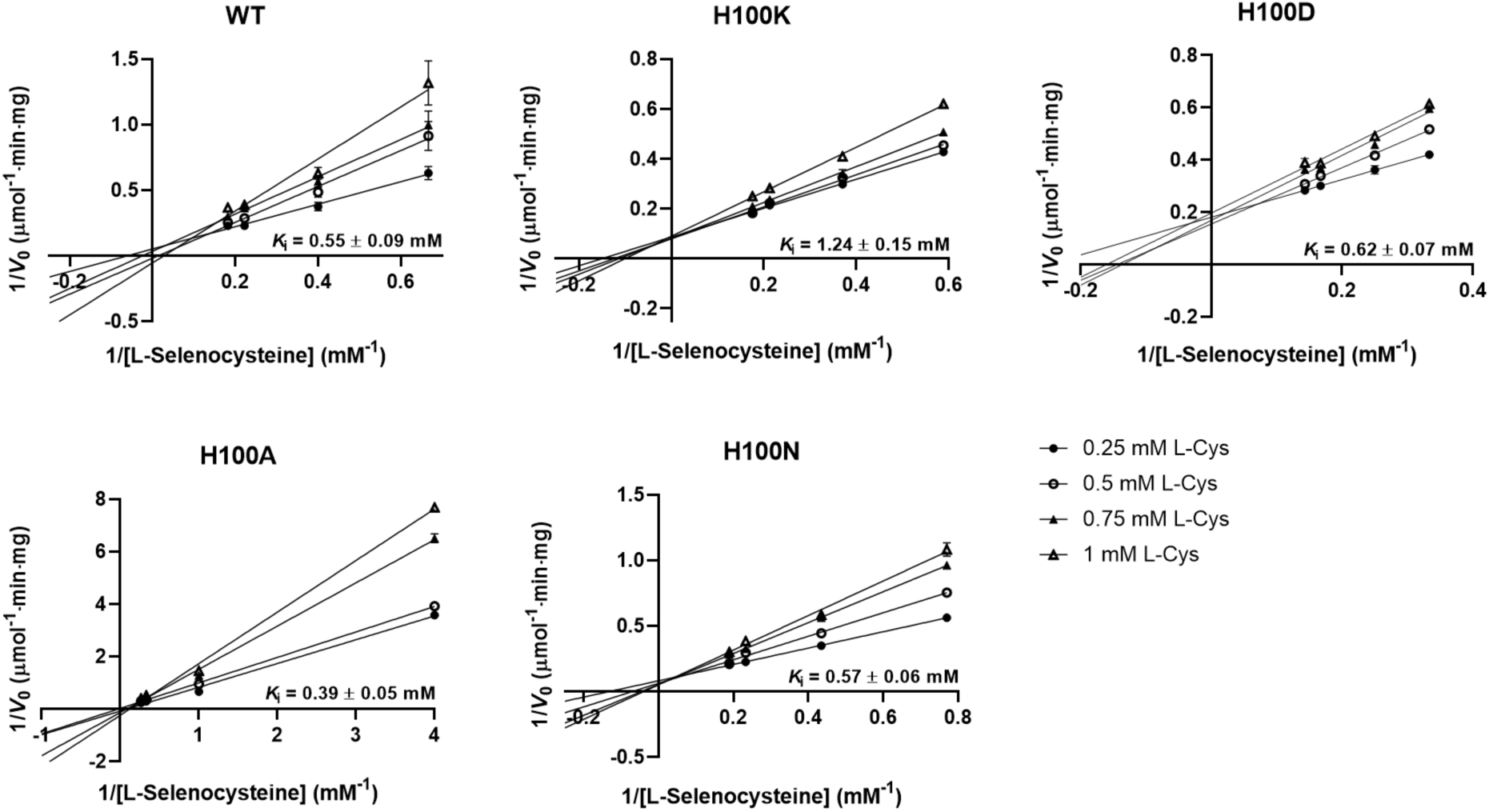
The selenocysteine lyase activities of SclA and H100 mutants are inhibited by L-cysteine in a competitive manner. The selenocysteine lyase activities of purified WT SclA and H100 mutants in the presence of increasing concentrations of L-cysteine were determined by the lead acetate assay. Four concentrations of L-selenocysteine (two above and below each *K*_m_^app^) were utilized. Double-reciprocal plots were generated from initial rates (*V*_0_) using the Lineweaver-Burk model in GraphPad Prism 8. *K*_i_^app^ and standard deviation values were calculated by fitting initial rates to the competitive inhibition model in the same program. Error bars indicate standard deviation; error bars smaller than the data points are not shown. In order to properly display the changes in kinetics between enzymes, the axes of each plot were not scaled equally.

To determine the *K*_i_^app^ values of each mutant, the same rates were fit with a competitive inhibition model in GraphPad Prism 8—the values and calculated error (standard deviation) are listed in Table 1. WT exhibited a *K*_i_^app^ of 0.55 ± 0.09 mM for L-cysteine while H100D and H100N exhibited similar values of 0.62 ± 0.07 mM and 0.57 ± mM (Fig. 4). Curiously enough, H100K demonstrated the weakest affinity for inhibitor with a *K*_i_^app^ of 1.24 ± 0.15 mM which accounted for a ∼2.3-fold increase compared to WT. On the other hand, H100A demonstrated the strongest affinity with a value of 0.39 ± 0.05 mM. Accounting for standard deviation, the *K*_i_^app^ for H100A was only slightly different from WT (∼1.4-fold decrease in *K*_i_^app^), so the actual significance of this value may be unclear. Strangely, all *K*_i_^app^ parameters were well below all reported *K*_m_^app^ values in this study, implying that L-cysteine binds substantially better to the enzyme compared to its primary substrate, L-selenocysteine. Overall, the variance in competitive inhibitory data further supports the notion that H100 influences catalysis and that mutagenesis of this residue perturbs the active site in a subtle manner.

## Discussion

In this work, we characterized the substrate specificity of SclA by investigating its enzyme kinetics after mutation of H100, a putative active site residue. For some enzymes, large changes in selenium recognition are due to a single point mutation. As stated previously, a D146K mutation of hSCL resulted in CDS activity (50,51). Likewise, a H240N variant of *Astragalus bisulcatus* CysRS was known to discriminate against selenocysteine (56). Moreover, using a bioinformatics and mutagenesis approach similar to this study, the authors found that orthologous histidine-to-asparagine CysRS mutants in *E. coli* and *Saccharomyces cerevisiae* resulted in decreased selenocysteine misincorporation and increased resistance to selenite toxicity (56).

Without a crystal structure of SclA, it is difficult to draw conclusions concerning the catalytic residues in its active site, but the alignment and homology models shown earlier can function as guides for analysis. It is also important to note that the p*K*_a_ of the catalytic cysteine, C341, is yet unknown. In water-based solutions, free cysteine has a p*K*_a_ of ∼8.5, resulting in mostly protonated and unreactive thiol groups at physiological pH. However, this is not always the case within the active site of a protein where catalytic cysteine residues are comparatively more reactive due to acidic p*K*_a_ values. The active site microenvironment of a protein contains clusters of differently-charged amino acid residues that greatly influence the p*K*_a_ values of catalytic cysteines—in this case, a lowering of the p*K*_a_ results in deprotonation to the reactive thiolate species at physiological pH. As stated earlier, CDSs contain a conserved reactive thiolate which non-specifically attacks sulfur or selenium, depending on the substrate (55). On the other hand, SCLs are thought to contain a thiol group rather than a thiolate, thereby limiting its reactivity to only L-selenocysteine rather than L-cysteine. It has been hypothesized that L-selenocysteine likely deprotonates the thiol to a thiolate which could then attack the electrophilic substrate—as stated earlier, L-cysteine does not exhibit this type of biochemical reactivity at physiological pH (51). In hSCL, D146 may have destabilized this thiolate group, restricting it to the protonated species (50,51). Thus, it would stand to reason that the specificity of SCLs most likely lies within controlling the protonation state of the catalytic cysteine via nearby residues.

In SclA, the function of H100 is still unclear. The significant changes in HSe^-^ kinetics from each mutant implies that it plays a role in catalysis; however, while the changes in kinetics amongst all enzymes were significant, there were only slight differences in turnover and catalytic efficiency (<2.0-fold differences in both *k*_cat_ and *k*_cat_/*K*_m_^app^). Therefore, H100 may not substantially influence catalysis as previously thought. Based on our modeling, it is more likely that this role may be played by H99 due to its close proximity to C341 (Fig. 2C). Recently, it has been found that H123 of *E. coli* SufS is required for CDS activity and that it may function as an acid-base catalyst in this mechanism (57). Moreover, the authors believed that H123 could deprotonate the catalytic thiol of C364 and share a π-π stacking interaction with PLP (57). Our sequence alignment (Fig. 1) revealed that H99 of SclA aligns with H123 of SufS; curiously, this histidine is found to be conserved in NifS-like proteins and is likely important for activity. Finally, the alignment reveals that this catalytic histidine is conserved as H145 in hSCL (Fig. 1). In the previous biochemical investigation of hSCL, the authors noted that H145 could possibly aid in determining substrate specificity by activating L-selenocysteine in the enzyme-substrate complex (51). Because SCLs are likely to contain a catalytic thiol rather than a thiolate, it is rather unlikely that H99 and H145 function as acid-base catalysts by deprotonating their respective catalytic cysteines. Instead, because the reactivity of L-selenocysteine may already achieve thiol deprotonation, it may be reasonable to suggest that these histidines help determine specificity for selenium by influencing the cysteine’s p*K*_a_. An investigation into the effects of H99 has not been attempted.

The inhibition of SCLs by L-cysteine is a puzzling phenomenon. Enzymes inhibited by a common metabolite present in high amounts would be extremely inefficient. In SclA, the *K*_i_^app^ for L-cysteine was determined to be 0.55 ± 0.09 mM while the *K*_m_^app^ for L-selenocysteine was 3.55 ± 0.32 mM. In hSCL, the *K*_i_^app^ is 5.85 mM while the *K*_m_^app^ for L-selenocysteine is 0.50 mM (34). In contrast to hSCL, SclA seems to have a greater affinity for L-cysteine than L-selenocysteine (∼6.5-fold difference between *K*_m_^app^ and *K*_i_^app^). The difference in binding affinities may suggest an overall distinction between the two enzymes— indeed, comparing prokaryotic and eukaryotic enzymes is difficult and may allow for misinterpretation of data without the aid of a crystal structure (in the case of SclA). In terms of kinetic changes, H100K exhibited the weakest affinity for L-cysteine as an inhibitor (∼2.3-fold increase in *K*_i_^app^). The lysyl residue may have changed the pH of the active site in a way that slightly disfavors binding of L-cysteine. Contrastingly, H100A had the highest binding affinity to L-cysteine, but it is unclear whether or not its *K*_i_^app^ is significant enough to make that claim (∼1.4-fold decrease). Overall, H100 mutagenesis did not seem to affect binding of L-cysteine as much as it did L-selenocysteine catalysis which may reveal something about the p*K*_a_ of C341’s sulfhydryl group.

Historically, thiol-based reductants like DTT have been used to assay the SCL and CDS activities of NifS-like enzymes, but this creates an inherent problem when determining the p*K*_a_ of the catalytic sulfhydryl: the reductant itself may be reacting with the sulfhydryl, thereby affecting the kinetics of the enzyme. Because DTT has been reported to nucleophilically attack and displace the sulfur atom bound to the catalytic thiolate in IscS, it can be assumed that a similar reaction probably occurs in SclA (28). In fact, it has been shown in Slr0387, a CDS in *Synechocystis* sp. PCC 6803, that the mechanism of persulfide cleavage by DTT is due to the dithiol forming a complex with PLP in order to position itself for attacking the catalytic sulfhydryl; moreover, the authors observed that this is the rate-determining step of catalysis, thus confounding any kinetic studies when DTT is the reductant (58). Furthermore, the authors made the observation that some CDSs including Slr0387 are subject to inhibition by L-cysteine when DTT is the reductant (58). They hypothesize that this may be due to both thiol-based molecules competing for the same catalytic sulfhydryl where, in one scenario, a DTT-PLP complex prevents L-cysteine from binding, and in another scenario, L-cysteine prevents DTT from binding to cofactor (58). Lastly, the authors found that using tris(2-carboxyethyl)phosphine (TCEP), a nonthiol, as a reductant was much more efficient in cleaving the persulfide bond compared to DTT (∼10-fold increase in *k*_cat_) and did not seem to exhibit a rate-limiting effect (58).

It is important to note that all *K*_m_^app^ values were in millimolar amounts, well above the physiological concentration of L-selenocysteine which is expected to be in low micromolar levels in the cell. Because HSe^-^ is incredibly toxic, uncontrolled release of this reactive species would not be favorable for any organism. Therefore, it may make sense that the relatively slow physiological turnover of this enzyme due to incredibly low substrate affinity serves as an internal control mechanism to prevent accidental poisoning of the cell. Moreover, the higher affinity for L-cysteine as evidenced by the *K*_i_^app^ values implies that the enzyme would almost always bind the inhibitor rather than its preferred substrate considering the fact that there is most likely ∼1000-fold more L-cysteine than L-selenocysteine in the cytosol. While this undoubtedly decreases the enzyme’s efficiency for L-selenocysteine degradation even more, one must further consider the repercussions of releasing free HSe^-^ into the cytosol such as thiol-mediated redox cycling (59,60). Coupled with its weak L-selenocysteine binding affinity, it is possible that the enzyme contains a sophisticated feedback mechanism to control HSe^-^ production as soon as it disengages the ribosome by sequestering itself with the large amount of cellular L-cysteine. If the enzyme were immediately “shut off” by the physiological concentration of L-cysteine to prevent HSe^-^ release, it may stand to reason that an additional factor may be required for the enzyme to “turn on” again and carry out its true biological function, whatever it may be.

While a role for SCLs has not been elucidated, it has been shown that some NifS-like enzymes can participate in mobilizing HSe^-^ from L-selenocysteine to SelD *in vitro* (61-63). The results of these mobilization studies heavily imply that the chemical species *in vivo* is most likely a selenium intermediate bound to the catalytic cysteine of SCLs to be transferred to another protein; in other words, SCLs probably do not randomly release HSe^-^ into the cellular environment. Because *E. faecalis* contains one known enzyme that requires selenium, it is hypothesized that SclA may contribute to the biosynthesis of that enzyme. Moreover, because this enzyme is a selenium-dependent XDH, *E. faecalis* therefore possesses this uncharacterized third trait of specific incorporation—thus, the generation of HSe^-^ from SclA may be the first step in this pathway. Indeed, the specificity of this enzyme towards L-selenocysteine would be crucial in synthesizing XDH.

In the same genetic locus as *sclA, selD*, and the gene encoding XDH, other genes have been identified as putatively involved in SDMH biosynthesis such as *yqeB* and *yqeC*. Past studies have shown that *yqeB* and *yqeC* generally co-localize with *selD* in the genome in many organisms lacking the ability to synthesize selenoproteins and selenium-containing tRNAs (44,48). These two genes may encode products that function to shuttle selenium in the SDMH biosynthetic pathway. The rationale for this “selenium-channeling” hypothesis stems from the fact that most CDSs are known to mobilize sulfur via protein-protein interactions; for example, in *A. vinelandii*, the Fe-S clusters in nitrogenase are synthesized by an interaction where NifS presents its thiol-bound sulfur intermediate to NifU, a scaffolding protein (64). In another example from both *A. vinelandii* and *E. coli*, IscS carries out a similar reaction with its partner, IscU (65,66). The similarities between the two subclasses of NifS-like proteins cannot be overstated—if CDSs can mobilize sulfur, surely SCLs can mobilize selenium. Because SCLs have low affinity for L-selenocysteine according to their millimolar *K*_m_^app^ constants, it is possible that they may require the presence of other proteins to vastly increase effective substrate affinity, assuming that they do mobilize selenium. Indeed, releasing HSe^-^ would be disastrous for the cell, and with the molecule itself being oxygen-labile, exposure to the cytosol would lead to oxidation to an unusable form. As of yet, SCLs are not known to have any *in vivo* electron acceptors, but the gene products of *selD, yqeB*, and *yqeC* may be likely candidates in the case of SclA. An analysis of SclA’s kinetics in the presence of all three proteins is recommended to probe this question.

## Experimental procedures

### Materials

Oligonucleotides were purchased from Integrated DNA Technologies. T4 DNA ligase, T4 DNA ligase reaction buffer, T4 Polynucleotide Kinase (PNK), Q5 High-Fidelity DNA Polymerase (Q5), Q5 reaction buffer, Q5 High GC Enhancer, Deoxynucleotide (dNTP) Solution Mix, and DpnI were all purchased from New England Biolabs. *E. coli* strains NEB 5-α and Lemo21(DE3) were purchased from New England Biolabs. Gel Loading Dye Blue (6×) and TriDye 1 kb DNA Ladder were purchased from New England Biolabs. Pierce Universal Nuclease for Cell Lysis and HisPur Cobalt Resin were purchased from Thermo Fisher Scientific. Spectra Multicolor Broad Range Protein Ladder was purchased from Thermo Fisher Scientific. The French Press and the Manual-Fill 40K Cell (FA-032) were purchased from Glen-Mills. Bio-Rad Protein Assay Dye Reagent Concentrate (Bradford reagent) was purchased from Bio-Rad. Q Sepharose Fast Flow was purchased from GE Healthcare. L-Selenocystine was purchased from Acros Organics. L-Cysteine hydrochloride monohydrate was purchased from Alfa Aesar.

### Bacterial strains and maintenance

All *E. coli* strains were maintained with lysogeny broth (LB) as per the Lennox formulation (1.0% tryptone, 0.5% yeast extract, 0.5% NaCl) (67-69). For solid media, LB was supplemented with 1.5% agar. Unless stated otherwise, *E. coli* strains used specifically for protein expression were maintained using LB supplemented with 2 KOH pellets per liter, 0.2% glucose, 0.2% glycerol, and 200 µg/mL ampicillin—this medium is referred to as “modified LB” in this work. Modified LB supplemented with 1.5% agar was used to maintain these strains on solid media. For overnight starter cultures, modified LB was supplemented with 1 mM MgSO_4_. For overexpression cultures, modified LB was supplemented with 1 mM MgSO_4_ and 0.1 mM pyridoxine, but glucose was omitted entirely. For starter cultures and smaller volumes, the Barnstead MaxQ 4000 orbital shaker was used. For overexpression cultures and larger volumes, the New Brunswick Innova 4230 shaker was used. All agitation steps were carried out at 240 rpm.

### Bioinformatic analysis

FASTA sequences of various characterized NifS-like enzymes were selected and aligned using Clustal Omega (70). These sequences were obtained from the following UniProt IDs: Q96I15 (hSCL), Q831E3 (*E. faecalis* SclA), O32164 (*B. subtilis* SufS), P77444 (*E. coli* SufS/CsdB), Q46925 (*E. coli* CSD/CsdA), P0A6B7 (*E. coli* IscS), and P05341 (*A. vinelandii* NifS) (71). The sequences were ordered by input as shown in Fig. 1.

To create 3-D structures of hSCL and SclA for active site analysis, the following methods were used. The Protein Data Bank (PDB) ID of hSCL, 3GZC, was used to generate a 3-D structure using PyMOL (72). Because no crystal structure of SclA existed at the time of this writing, it had to be modeled in a different manner. The aforementioned FASTA sequence of SclA was used to find templates of closest putative structural homology using ExPASy’s SWISS-MODEL (73). The SclA sequence was then threaded through the top result, *B. subtilis* SufS (PDB ID: 5J8Q), to generate a hypothetical structural model. The data for this homology model were converted to a PDB format and then used to generate a 3-D structure using PyMOL. D146K and H100 mutants were also modeled with the same software.

### Cloning and mutagenesis

The pET100 vector designated as pRT1 containing *sclA* (EF2568) cloned from *E. faecalis* V583 was subjected to ‘Round-the-horn site-directed mutagenesis (53,54). Briefly, primers were designed with the intent to mutate the histidine at amino acid position 100 to the following residues: lysine (H100K), aspartate (H100D), alanine (H100A), and asparagine (H100N). Forward primers contained the mutated codons on the 5’ ends while the reverse primer was designed to bind directly next to the site of mutation. All primers were designed to have a melting temperature of ∼60 °C and are referenced in Table S1. Primers were phosphorylated by incubating a 50 µL mixture containing 1× T4 DNA ligase reaction buffer, 10 µM primers, and 1 µL of PNK (10,000 U/mL) at 37 °C for 1 hour; afterwards, the PNK was inactivated with a final 5-minute incubation at 95 °C. For each point mutant, several 50 µL master mixes of the following reagents were prepared on ice: 1× Q5 reaction buffer, 1× Q5 High GC Enhancer, 300 nM forward and reverse primers, 0.2 mM dNTPs, 0.5 ng of pRT1, and 0.5 µL of Q5 (2,000 U/mL). To perform the polymerase chain reaction (PCR), the master mixes were placed in a Techne TC-312 thermal cycler with a pre-heated lid and were initially denatured at 98 °C for 1 minute. Afterwards, the following program was cycled through 27 times: denaturation at 96 °C for 30 seconds, annealing at 64 °C for 30 seconds, and extension at 72 °C for 14 minutes. Finally, samples were cooled at 18 °C for 1 minute. To check for PCR products, 5 µL samples were mixed with 2 µL of Gel Loading Dye Blue (6×). From these samples, 5 µL along with 5 µL of TriDye 1 kb DNA Ladder were loaded on a 1% agarose gel prepared with 1× Tris-acetate-EDTA (TAE, 40 mM tris(hydroxymethyl)aminomethane [Tris], 20 mM acetate, 10 mM ethylenediaminetetraacetate [EDTA]) and 0.5 µg/mL ethidium bromide and were electrophoresed at 100 V for 30 minutes.

After mutagenesis, samples were treated with 1 µL of DpnI (20,000 U/mL) and incubated at 37 °C for 1 hour to digest the template plasmid. Digested samples were electrophoresed on a 1% agarose gel prepared with 1× TAE and 0.005% crystal violet at 100 V for 30 minutes. The target bands were excised with a sterile razor blade, transferred to 1.7 mL microcentrifuge tubes containing 500 µL of gel melting buffer (5.5 M guanidinium thiocyanate, 0.1 M sodium acetate, [pH 5.0]), and melted in a Fisher Isotemp 210 water bath at 50 °C for 10 minutes. After samples were fully dissolved and denatured, 150 µL of 100% isopropanol were added to each tube and vortexed thoroughly. Unless otherwise stated, all centrifugation steps regarding spin columns for the rest of this work were carried out using an Eppendorf 5418 centrifuge. Denatured samples were bound to DNA Mini-Spin Columns (Enzymax LLC) and centrifuged twice at 2,000 × g for 1 minute to discard the non-bound flow-through. An additional 300 µL of gel melting buffer were added to each sample, and each tube was centrifuged at 2,000 × g for 1 minute. Samples were washed with a column wash buffer (5 mM 4-(2-hydroxyethyl)-1-piperazineethanesulfonate-KOH [HEPES-KOH], 20 mM NaCl, 0.02 mM EDTA, 80% ethanol, [pH 8.3]) three times at decreasing volumes per wash: 400 µL for the first wash, 250 µL for the second wash, and 50 µL for the final wash—all samples were centrifuged at 2,000 × g for 1 minute for every subsequent wash to remove the buffer. Afterwards, the samples were centrifuged at 16,900 × g for 1 minute to remove any residual ethanol. Finally, 50 µL of column elution buffer (2.5 mM Tris-HCl, 0.1 mM EDTA, [pH 8.0]) were transferred to the center of each column; purified products were allowed to incubate at room temperature for 1 minute before finally being eluted by centrifugation at 16,900 × g for 1 minute.

Purified products were ligated by incubating a 10 µL mixture containing 1× T4 DNA ligase reaction buffer, 8 µL of PCR product, and 0.4 µL of T4 DNA Ligase (400,000 U/mL) at room temperature for 10 minutes. After ligation, mutant plasmids were transformed into *E. coli* strain NEB 5-α. Briefly, NEB 5-α was streaked onto an LB plate and incubated overnight at 37 °C. A single colony was selected and used to inoculate a 1 mL starter culture composed of LB and 0.2% glucose which aerobically grew overnight at 37 °C. Starter cultures were diluted 1:100 into several 1 mL broths containing LB and 0.2% glucose and allowed to grow for 2 hours at 37 °C. After reaching mid-exponential phase, cultures were harvested at 5,000 × g for 3 minutes. After discarding media, pellets were resuspended in 100 µL Transformation and Storage Solution (TSS, 10% polyethylene glycol 8000, 30 mM MgCl_2_, 5% dimethyl sulfoxide, 0.2% glucose, dissolved in LB), and 40 µL of resuspended cells were transferred to pre-chilled 1.7 mL microcentrifuge tubes (74). Five microliters of each ligated PCR product were transferred to their respective 40 µL mixtures to begin transformation. Competent cells were incubated on ice for 10 minutes, at room temperature for 10 minutes, and finally on ice again for an additional 10 minutes to allow for transformation of ligated products. One milliliter of LB with 0.2% glucose was added to each transformant tube, and all tubes were incubated at 37 °C for 1 hour to allow for cell recovery. Recovered cells were harvested at 5,000 × g for 3 minutes, and ∼900 µL of supernatant were discarded. Pellets were resuspended with the remaining ∼100 µL of supernatant and transferred to LB agar plates supplemented with 100 µg/mL ampicillin. Plates were incubated overnight at 37 °C. Several transformant colonies were selected for minipreps via inoculation of several 2 mL LB broths which were cultured overnight at 37 °C. Plasmid DNA was extracted and purified from each culture via spin-columns. Mutations were confirmed by Sanger sequencing provided by GENEWIZ—chromatograms were generated using the sequencing data and visibly inspected using SnapGene Viewer (GSL Biotech).

### Protein overexpression

WT SclA and mutants were overexpressed in *E. coli* strain Lemo21(DE3) and purified using N-terminal His_6_-tags. Unless stated otherwise, the following procedure was used for overexpression and purification of WT and all mutants mentioned in this study. Glycerol stocks of these strains were streaked onto modified LB agar and incubated overnight at 37 °C. Pure colonies were used to inoculate 50 mL starter cultures which were allowed to shake overnight at 37 °C. Starter cultures were diluted 1:100 into 4 liter (2 × 2 liter) overexpression cultures which were allowed to grow for 1.5 hours at 37 °C. The temperature was then immediately shifted to 25 °C and allowed to equilibrate for an additional 20 minutes. After equilibration, the cultures were induced with 0.5 mM isopropyl β-D-1-thiogalactopyranoside (IPTG) for 7 hours. Cultures were immediately placed on ice, transferred to pre-chilled 1-liter centrifuge bottles, and harvested at 12,000 × g for 10 minutes at 4 °C using a Sorvall RC6+ centrifuge—this device was used for all harvest and clarification steps unless otherwise stated. Cell pellets were resuspended with 25 mL of cold cell wash buffer (25 mM Tris-HCl, 100 mM NaCl, 1 mM EDTA, [pH 8.0]), harvested as stated before, and stored at −80 °C.

### Protein purification

Before lysis, cells were thawed at 4 °C, resuspended in 80 mL of cold lysis buffer (25 mM Tris-HCl, 300 mM NaCl, 0.05% Tween-20, 5% glycerol, [pH 8.0]) supplemented with 1 mM benzamidine, and vortexed for 1 minute. One microliter of Pierce Universal Nuclease for Cell Lysis (250 U/µL) was added to the diluted suspension to digest any remaining nucleic acids. The suspension was passed through a French Press with a pre-chilled Manual-Fill 40K Cell three times at maximum standard pressure (40,000 psi) to completely lyse the cells. All centrifugation and purification steps were carried out at 4 °C. The crude lysate was clarified at 28,900 × g for 20 minutes. From this point onward, all centrifugation steps regarding resin harvest were carried out using a Hermle Z400K centrifuge. Two milliliters of HisPur Cobalt Resin were washed with 20 mL of cobalt binding buffer (25 mM Tris-HCl, 300 mM NaCl, 5 mM imidazole, 0.05% Tween-20, 14 mM β-mercaptoethanol [β-Me], 5% glycerol, [pH 8.0]) and centrifuged at 1,000 × g for 5 minutes. After decanting the supernatant, the washed resin was mixed with the soluble lysate in a 50 mL conical tube and allowed to batch-bind via end over end rotation for 1 hour at 4 °C. The mixture was centrifuged at 1,000 × g for 5 minutes to collect the protein-bound resin. From here on out, the Bradford assay was utilized to track the protein across the purification by sampling 5 µL from each step and adding to 100 µL of 1× Bradford reagent in a 96-well plate (75). Resin was resuspended with cobalt binding buffer, transferred to a gravity column, and pre-washed with cobalt wash buffer (25 mM Tris-HCl, 300 mM NaCl, 20 mM imidazole, 0.05% Tween-20, 14 mM β-Me, 5% glycerol, [pH 8.0]) at 75% of the bed volume. Afterwards, the resin was washed with 3 bed volumes of cobalt wash buffer at least 4 times. To dilute the wash buffer, the resin was then pre-eluted with cobalt elution buffer (25 mM Tris-HCl, 300 mM NaCl, 250 mM imidazole, 0.05% Tween-20, 14 mM β-Me, 5% glycerol, [pH 8.0]) at 75% of the bed volume. Afterwards, the target protein was eluted with 1 bed volume of cobalt elution buffer at least 4 times. Once cobalt affinity was completed, 1 mL of Q Sepharose Fast Flow resin was washed with 20 mL of Q binding buffer (12.5 mM Tris-HCl, 50 mM NaCl, 0.05% Tween-20, 14 mM β-Me, 5% glycerol, 0.1 mM EDTA, [pH 8.0]). Elutions were pooled together and mixed with Q resin; to dilute the imidazole, the total volume of the binding mixture was brought up to 40 mL with deionized water (dH_2_O). Protein was allowed to bind for 15 minutes in the cold as described earlier. Protein-bound resin was harvested via centrifugation at 1,000 × g for 5 minutes. Resin was resuspended with Q binding buffer, transferred to a gravity column, and pre-washed with Q wash buffer (12.5 mM Tris-HCl, 100 mM NaCl, 0.05% Tween-20, 14 mM β-Me, 5% glycerol, 0.1 mM EDTA, [pH 8.0]) at 75% of the bed volume. Two bed volumes of Q wash buffer were used to wash the resin at least 2 times. To dilute the wash buffer, the resin was pre-eluted with Q elution buffer (12.5 mM Tris-HCl, 500 mM NaCl, 0.05% Tween-20, 14 mM β-Me, 5% glycerol, 0.1 mM EDTA, [pH 8.0]) at 75% of the bed volume. Afterwards, protein was eluted with 1 bed volume of Q elution buffer at least 6 times. Elutions were pooled together and dialyzed against 1 liter of storage buffer (50 mM HEPES-KOH, 5% glycerol, [pH 7.5]) overnight at 4 °C.

### SDS-PAGE analysis

Induction and purification of all proteins were analyzed via sodium dodecyl sulfate-polyacrylamide gel electrophoresis (SDS-PAGE) (76). To assess the induction of each protein, 1 mL samples of the overexpression cultures were taken before and after full protein induction. The sample before addition of IPTG was labeled “pre-IPTG” while the sample after full induction was labeled “post-IPTG”. The optical densities at 600 nm (OD_600_) of each sample were analyzed using an Agilent 8453 UV-visible spectrophotometer. The OD_600_ of the post-IPTG sample was normalized to the OD_600_ of the pre-IPTG sample. Both samples were harvested at 12,000 × g for 5 minutes using an Eppendorf 5418 centrifuge. After decanting media, pellets were stored at −80 °C. For sample preparation of induction pellets, 40 µL of 2× Laemmli sample buffer were used to resuspend the cells before lysing via incubation at 100 °C in a sand bath for 5 minutes. For general sample preparation, 8 µL were pulled from all steps of the purification scheme, mixed with 32 µL of 2× Laemmli sample buffer, and heated similarly for 5 minutes. Five microliters of boiled samples were routinely loaded on 12% Tris-glycine gels and electrophoresed at 35 mA for 1 hour. Gels were gently washed with dH_2_O for 5 minutes before replacing the water with a gel staining solution (0.02% Coomassie Brilliant Blue G-250, 10% methanol, 10% acetate) and microwaving until the staining solution began to boil. After initial staining, gels were agitated on a low-speed orbital shaker for 10 minutes before replacing staining solution with 10% acetate for destaining. To accelerate the destaining process, the solution was microwaved to a boil, and Kimwipes were added during agitation. Gels with clear backgrounds were deemed ready for imaging. Destained gels were gently washed with dH_2_O before being analyzed with a ChemiDoc MP Imaging System.

### Protein quantification

Protein concentrations of dialyzed enzymes were quantified via their absorbance profiles at 280 nm. Briefly, samples were diluted 1:10 in 6 M guanidinium-HCl and read using an Agilent 8453 UV-visible spectrophotometer. The predicted molecular weight and theoretical molar absorptivity coefficient of SclA at 280 nm were 41,525.86 Da and 39,420 M^-1^ cm^-1^ respectively, as determined by the ProtParam tool by ExPASy (77). Beer’s law was used to calculate protein concentration (78). Proteins were stored in 400 µL aliquots at −20 °C.

### Lead acetate assay

The SCL activities of all purified proteins were analyzed with the lead acetate assay (32). Upon addition of lead acetate, any hydrogen selenide (H_2_Se) present reacts to form lead selenide (PbSe) which can be detected at 400 nm. The assay was carried out in a 96-well plate format. The assay mixture was formulated containing the following reagents: 50 mM HEPES-KOH (pH 7.5), 5 mM DTT, 0.2 mM PLP, 1 mM pyruvate, and 0.02 mM EDTA. The substrate was prepared by first dissolving the oxidized form, L-selenocystine, with NaOH at a 1:1 molar ratio of L-selenocysteine-to-NaOH—the diselenide molecule is automatically reduced by DTT to L-selenocysteine in the assay. An enzyme mixture was prepared containing 0.3 mg/mL SclA and 0.8 mg/mL bovine serum albumin (BSA); from this mix, 5 µL were added to 90 µL of 1× assay mixture. To begin the enzymatic reaction, 5 µL of prepared L-selenocystine were added to the above mixture for a final volume of 100 µL. The assay was carried out in triplicate at 25 °C for various lengths of time to ascertain the initial rates of HSe^-^ production at varying substrate concentrations. Enzyme was omitted in a blank as a negative control. Reactions were quenched with 100 µL of 5 mM lead acetate dissolved in 0.5 M HCl. The optical densities at 400 nm of 200 µL quenched samples were analyzed with a SpectraMax 190 Microplate Reader. The molar turbidity coefficient of PbSe at 400 nm is 1.18 × 10^4^ M^-1^ cm^-1^ and was used to calculate the HSe^-^ concentration via Beer’s law (30,78). Blank rates were averaged and subtracted from sample rates. Rates were fit to the Michaelis-Menten equation using GraphPad Prism 8. For inhibition studies, several solutions of 1× assay mixture were supplemented with 0.25 mM, 0.5 mM, 0.75 mM, and 1 mM L-cysteine. For inhibition assays, rates were transformed to generate double-reciprocal plots; after inspection of the trendlines in these plots, rates were fit to a competitive inhibition model using GraphPad Prism 8. All kinetic parameters and standard deviations were calculated using GraphPad Prism 8.

## Acknowledgments

The authors would like to thank Rebecca Tarrien for contributing to this work, specifically in cloning the *sclA* gene from *Enterococcus faecalis* V583 genomic DNA into pET100/D-TOPO to enable these biochemical analyses.

## Conflict of interest

No conflicts from any authors.

## Author contributions

M.A.J. and W.T.S. designed the experiments, S.J.N. and C.V.G. carried out experiments on SclA in their thesis work that led to these experiments, M.A.J. drafted and edited the manuscript and W.T.S. edited the final version of the manuscript. All authors reviewed and edited the manuscript prior to submission.

## FOOTNOTES

W.T.S. and S.J.N. acknowledge support from the NIH NIAID 1R21AI101560-01 to contributing to this work.

## References

1. Cone, J. E., Del Rio, R. M., Davis, J. N., and Stadtman, T. C. (1976) Chemical characterization of the selenoprotein component of clostridial glycine reductase: identification of selenocysteine as the organoselenium moiety. Proc Natl Acad Sci U S A 73, 2659–2663

2. Bock, A., Forchhammer, K., Heider, J., Leinfelder, W., Sawers, G., Veprek, B., and Zinoni, F. (1991) Selenocysteine: the 21st amino acid. Mol Microbiol 5, 515–520

3. Berry, M. J., Banu, L., Chen, Y. Y., Mandel, S. J., Kieffer, J. D., Harney, J. W., and Larsen, P. R. (1991) Recognition of UGA as a selenocysteine codon in type I deiodinase requires sequences in the 3’ untranslated region. Nature 353, 273–276

4. Gonzalez-Flores, J. N., Shetty, S. P., Dubey, A., and Copeland, P. R. (2013) The molecular biology of selenocysteine. Biomol Concepts 4, 349–365

5. Leinfelder, W., Forchhammer, K., Zinoni, F., Sawers, G., Mandrand-Berthelot, M. A., and Bock, A. (1988) Escherichia coli genes whose products are involved in selenium metabolism. J Bacteriol 170, 540–546

6. Sawers, G., Heider, J., Zehelein, E., and Bock, A. (1991) Expression and operon structure of the sel genes of Escherichia coli and identification of a third selenium-containing formate dehydrogenase isoenzyme. J Bacteriol 173, 4983–4993

7. Huber, R. E., and Criddle, R. S. (1967) Comparison of the chemical properties of selenocysteine and selenocystine with their sulfur analogs. Arch Biochem Biophys 122, 164–173

8. Snider, G. W., Ruggles, E., Khan, N., and Hondal, R. J. (2013) Selenocysteine confers resistance to inactivation by oxidation in thioredoxin reductase: comparison of selenium and sulfur enzymes. Biochemistry 52, 5472–5481

9. Axley, M. J., Bock, A., and Stadtman, T. C. (1991) Catalytic properties of an Escherichia coli formate dehydrogenase mutant in which sulfur replaces selenium. Proc Natl Acad Sci U S A 88, 8450–8454

10. Berry, M. J., Banu, L., and Larsen, P. R. (1991) Type I iodothyronine deiodinase is a selenocysteine-containing enzyme. Nature 349, 438–440

11. Kim, M. J., Lee, B. C., Hwang, K. Y., Gladyshev, V. N., and Kim, H. Y. (2015) Selenium utilization in thioredoxin and catalytic advantage provided by selenocysteine. Biochem Biophys Res Commun 461, 648–652

12. Ching, W. M., Alzner-DeWeerd, B., and Stadtman, T. C. (1985) A selenium-containing nucleoside at the first position of the anticodon in seleno-tRNAGlu from Clostridium sticklandii. Proc Natl Acad Sci U S A 82, 347–350

13. Wolfe, M. D., Ahmed, F., Lacourciere, G. M., Lauhon, C. T., Stadtman, T. C., and Larson, T. J. (2004) Functional diversity of the rhodanese homology domain: the Escherichia coli ybbB gene encodes a selenophosphate-dependent tRNA 2-selenouridine synthase. J Biol Chem 279, 1801–1809

14. Wittwer, A. J. (1983) Specific incorporation of selenium into lysine- and glutamate-accepting tRNAs from Escherichia coli. J Biol Chem 258, 8637–8641

15. Wagner, R., and Andreesen, J. R. (1979) Selenium requirement for active xanthine dehydrogenase from Clostridium acidiurici and Clostridium cylindrosporum. Arch Microbiol 121, 255–260

16. Self, W. T., Wolfe, M. D., and Stadtman, T. C. (2003) Cofactor determination and spectroscopic characterization of the selenium-dependent purine hydroxylase from Clostridium purinolyticum. Biochemistry 42, 11382–11390

17. Self, W. T., and Stadtman, T. C. (2000) Selenium-dependent metabolism of purines: A selenium-dependent purine hydroxylase and xanthine dehydrogenase were purified from Clostridium purinolyticum and characterized. Proc Natl Acad Sci U S A 97, 7208–7213

18. Gladyshev, V. N., Khangulov, S. V., and Stadtman, T. C. (1994) Nicotinic acid hydroxylase from Clostridium barkeri: electron paramagnetic resonance studies show that selenium is coordinated with molybdenum in the catalytically active selenium-dependent enzyme. Proc Natl Acad Sci U S A 91, 232–236

19. Muller, S., Heider, J., and Bock, A. (1997) The path of unspecific incorporation of selenium in Escherichia coli. Arch Microbiol 168, 421–427

20. Young, P. A., and Kaiser, II. (1975) Aminoacylation of Escherichia coli cysteine tRNA by selenocysteine. Arch Biochem Biophys 171, 483–489

21. Zorn, M., Ihling, C. H., Golbik, R., Sawers, R. G., and Sinz, A. (2013) Selective selC-independent selenocysteine incorporation into formate dehydrogenases. PLoS One 8, e61913

22. Brown, T. A., and Shrift, A. (1980) Identification of Selenocysteine in the Proteins of Selenate-grown Vigna radiata. Plant Physiol 66, 758–761

23. Eustice, D. C., Foster, I., Kull, F. J., and Shrift, A. (1980) In Vitro Incorporation of Selenomethionine into Protein by Vigna radiata Polysomes. Plant Physiol 66, 182–186

24. Eustice, D. C., Kull, F. J., and Shrift, A. (1981) In vitro incorporation of selenomethionine into protein by astragalus polysomes. Plant Physiol 67, 1059–1060

25. Burk, R. F., Hill, K. E., and Motley, A. K. (2001) Plasma selenium in specific and non-specific forms. Biofactors 14, 107–114

26. Zheng, L., White, R. H., Cash, V. L., Jack, R. F., and Dean, D. R. (1993) Cysteine desulfurase activity indicates a role for NIFS in metallocluster biosynthesis. Proc Natl Acad Sci U S A 90, 2754–2758

27. Mihara, H., and Esaki, N. (2002) Bacterial cysteine desulfurases: their function and mechanisms. Appl Microbiol Biotechnol 60, 12–23

28. Flint, D. H. (1996) Escherichia coli contains a protein that is homologous in function and N-terminal sequence to the protein encoded by the nifS gene of Azotobacter vinelandii and that can participate in the synthesis of the Fe-S cluster of dihydroxy-acid dehydratase. J Biol Chem 271, 16068–16074

29. Mihara, H., Kurihara, T., Yoshimura, T., Soda, K., and Esaki, N. (1997) Cysteine sulfinate desulfinase, a NIFS-like protein of Escherichia coli with selenocysteine lyase and cysteine desulfurase activities. Gene cloning, purification, and characterization of a novel pyridoxal enzyme. J Biol Chem 272, 22417–22424

30. Mihara, H., Maeda, M., Fujii, T., Kurihara, T., Hata, Y., and Esaki, N. (1999) A nifS-like gene, csdB, encodes an Escherichia coli counterpart of mammalian selenocysteine lyase. Gene cloning, purification, characterization and preliminary x-ray crystallographic studies. J Biol Chem 274, 14768–14772

31. Mihara, H., Kurihara, T., Yoshimura, T., and Esaki, N. (2000) Kinetic and mutational studies of three NifS homologs from Escherichia coli: mechanistic difference between L-cysteine desulfurase and L-selenocysteine lyase reactions. J Biochem 127, 559–567

32. Esaki, N., Nakamura, T., Tanaka, H., and Soda, K. (1982) Selenocysteine lyase, a novel enzyme that specifically acts on selenocysteine. Mammalian distribution and purification and properties of pig liver enzyme. J Biol Chem 257, 4386–4391

33. Chocat, P., Esaki, N., Tanizawa, K., Nakamura, K., Tanaka, H., and Soda, K. (1985) Purification and characterization of selenocysteine beta-lyase from Citrobacter freundii. J Bacteriol 163, 669–676

34. Daher, R., and Van Lente, F. (1992) Characterization of selenocysteine lyase in human tissues and its relationship to tissue selenium concentrations. J Trace Elem Electrolytes Health Dis 6, 189–194

35. Veres, Z., Kim, I. Y., Scholz, T. D., and Stadtman, T. C. (1994) Selenophosphate synthetase. Enzyme properties and catalytic reaction. J Biol Chem 269, 10597–10603

36. Xu, X. M., Carlson, B. A., Irons, R., Mix, H., Zhong, N., Gladyshev, V. N., and Hatfield, D. L. (2007) Selenophosphate synthetase 2 is essential for selenoprotein biosynthesis. Biochem J 404, 115–120

37. Kurokawa, S., Takehashi, M., Tanaka, H., Mihara, H., Kurihara, T., Tanaka, S., Hill, K., Burk, R., and Esaki, N. (2011) Mammalian selenocysteine lyase is involved in selenoprotein biosynthesis. J Nutr Sci Vitaminol (Tokyo) 57, 298–305

38. Tobe, R., Mihara, H., Kurihara, T., and Esaki, N. (2009) Identification of proteins interacting with selenocysteine lyase. Biosci Biotechnol Biochem 73, 1230–1232

39. Seale, L. A., Hashimoto, A. C., Kurokawa, S., Gilman, C. L., Seyedali, A., Bellinger, F. P., Raman, A. V., and Berry, M. J. (2012) Disruption of the selenocysteine lyase-mediated selenium recycling pathway leads to metabolic syndrome in mice. Mol Cell Biol 32, 4141–4154

40. Seale, L. A., Gilman, C. L., Hashimoto, A. C., Ogawa-Wong, A. N., and Berry, M. J. (2015) Diet-induced obesity in the selenocysteine lyase knockout mouse. Antioxid Redox Signal 23, 761–774

41. Seale, L. A., Khadka, V. S., Menor, M., Xie, G., Watanabe, L. M., Sasuclark, A., Guirguis, K., Ha, H. Y., Hashimoto, A. C., Peplowska, K., Tiirikainen, M., Jia, W., Berry, M. J., and Deng, Y. (2019) Combined Omics Reveals That Disruption of the Selenocysteine Lyase Gene Affects Amino Acid Pathways in Mice. Nutrients 11

42. Mihara, H., Kurihara, T., Watanabe, T., Yoshimura, T., and Esaki, N. (2000) cDNA cloning, purification, and characterization of mouse liver selenocysteine lyase. Candidate for selenium delivery protein in selenoprotein synthesis. J Biol Chem 275, 6195–6200

43. Chocat, P., Esaki, N., Nakamura, T., Tanaka, H., and Soda, K. (1983) Microbial distribution of selenocysteine lyase. J Bacteriol 156, 455–457

44. Haft, D. H., and Self, W. T. (2008) Orphan SelD proteins and selenium-dependent molybdenum hydroxylases. Biol Direct 3, 4

45. Agudelo Higuita, N. I., and Huycke, M. M. (2014) Enterococcal Disease, Epidemiology, and Implications for Treatment. in Enterococci: From Commensals to Leading Causes of Drug Resistant Infection (Gilmore, M. S., Clewell, D. B., Ike, Y., and Shankar, N. eds.), Boston. pp

46. Mohamed, J. A., and Huang, D. B. (2007) Biofilm formation by enterococci. J Med Microbiol 56, 1581–1588

47. Zhang, Y., Romero, H., Salinas, G., and Gladyshev, V. N. (2006) Dynamic evolution of selenocysteine utilization in bacteria: a balance between selenoprotein loss and evolution of selenocysteine from redox active cysteine residues. Genome Biol 7, R94

48. Zhang, Y., Turanov, A. A., Hatfield, D. L., and Gladyshev, V. N. (2008) In silico identification of genes involved in selenium metabolism: evidence for a third selenium utilization trait. BMC Genomics 9, 251

49. Srivastava, M., Mallard, C., Barke, T., Hancock, L. E., and Self, W. T. (2011) A selenium-dependent xanthine dehydrogenase triggers biofilm proliferation in Enterococcus faecalis through oxidant production. J Bacteriol 193, 1643–1652

50. Collins, R., Johansson, A. L., Karlberg, T., Markova, N., van den Berg, S., Olesen, K., Hammarstrom, M., Flores, A., Schuler, H., Schiavone, L. H., Brzezinski, P., Arner, E. S., and Hogbom, M. (2012) Biochemical discrimination between selenium and sulfur 1: a single residue provides selenium specificity to human selenocysteine lyase. PLoS One 7, e30581

51. Johansson, A. L., Collins, R., Arner, E. S., Brzezinski, P., and Hogbom, M. (2012) Biochemical discrimination between selenium and sulfur 2: mechanistic investigation of the selenium specificity of human selenocysteine lyase. PLoS One 7, e30528

52. Blauenburg, B., Mielcarek, A., Altegoer, F., Fage, C. D., Linne, U., Bange, G., and Marahiel, M. A. (2016) Crystal Structure of Bacillus subtilis Cysteine Desulfurase SufS and Its Dynamic Interaction with Frataxin and Scaffold Protein SufU. PLoS One 11, e0158749

53. Moore, S. D. (2018) ’Round-the-horn site-directed mutagenesis. in OpenWetWare

54. Moore, S. D., and Prevelige, P. E., Jr. (2002) A P22 scaffold protein mutation increases the robustness of head assembly in the presence of excess portal protein. J Virol 76, 10245–10255

55. Zheng, L., White, R. H., Cash, V. L., and Dean, D. R. (1994) Mechanism for the desulfurization of L-cysteine catalyzed by the nifS gene product. Biochemistry 33, 4714–4720

56. Hoffman, K. S., Vargas-Rodriguez, O., Bak, D. W., Mukai, T., Woodward, L. K., Weerapana, E., Soll, D., and Reynolds, N. M. (2019) A cysteinyl-tRNA synthetase variant confers resistance against selenite toxicity and decreases selenocysteine misincorporation. J Biol Chem 294, 12855–12865

57. Blahut, M., Wise, C. E., Bruno, M. R., Dong, G., Makris, T. M., Frantom, P. A., Dunkle, J. A., and Outten, F. W. (2019) Direct observation of intermediates in the SufS cysteine desulfurase reaction reveals functional roles of conserved active-site residues. J Biol Chem 294, 12444–12458

58. Behshad, E., Parkin, S. E., and Bollinger, J. M., Jr. (2004) Mechanism of cysteine desulfurase Slr0387 from Synechocystis sp. PCC 6803: kinetic analysis of cleavage of the persulfide intermediate by chemical reductants. Biochemistry 43, 12220–12226

59. Spallholz, J. E. (1997) Free radical generation by selenium compounds and their prooxidant toxicity. Biomed Environ Sci 10, 260–270

60. Kumar, S., Bjornstedt, M., and Holmgren, A. (1992) Selenite is a substrate for calf thymus thioredoxin reductase and thioredoxin and elicits a large non-stoichiometric oxidation of NADPH in the presence of oxygen. Eur J Biochem 207, 435–439

61. Lacourciere, G. M., and Stadtman, T. C. (1998) The NIFS protein can function as a selenide delivery protein in the biosynthesis of selenophosphate. J Biol Chem 273, 30921–30926

62. Lacourciere, G. M., Mihara, H., Kurihara, T., Esaki, N., and Stadtman, T. C. (2000) Escherichia coli NifS-like proteins provide selenium in the pathway for the biosynthesis of selenophosphate. J Biol Chem 275, 23769–23773

63. Lacourciere, G. M. (2002) Selenium is mobilized in vivo from free selenocysteine and is incorporated specifically into formate dehydrogenase H and tRNA nucleosides. J Bacteriol 184, 1940–1946

64. Yuvaniyama, P., Agar, J. N., Cash, V. L., Johnson, M. K., and Dean, D. R. (2000) NifS-directed assembly of a transient [2Fe-2S] cluster within the NifU protein. Proc Natl Acad Sci U S A 97, 599–604

65. Urbina, H. D., Silberg, J. J., Hoff, K. G., and Vickery, L. E. (2001) Transfer of sulfur from IscS to IscU during Fe/S cluster assembly. J Biol Chem 276, 44521–44526

66. Smith, A. D., Agar, J. N., Johnson, K. A., Frazzon, J., Amster, I. J., Dean, D. R., and Johnson, M. K. (2001) Sulfur transfer from IscS to IscU: the first step in iron-sulfur cluster biosynthesis. J Am Chem Soc 123, 11103–11104

67. Bertani, G. (1951) Studies on lysogenesis. I. The mode of phage liberation by lysogenic Escherichia coli. J Bacteriol 62, 293–300

68. Lennox, E. S. (1955) Transduction of linked genetic characters of the host by bacteriophage P1. Virology 1, 190–206

69. Bertani, G. (2004) Lysogeny at mid-twentieth century: P1, P2, and other experimental systems. J Bacteriol 186, 595–600

70. Sievers, F., Wilm, A., Dineen, D., Gibson, T. J., Karplus, K., Li, W., Lopez, R., McWilliam, H., Remmert, M., Soding, J., Thompson, J. D., and Higgins, D. G. (2011) Fast, scalable generation of high-quality protein multiple sequence alignments using Clustal Omega. Mol Syst Biol 7, 539

71. UniProt, C. (2019) UniProt: a worldwide hub of protein knowledge. Nucleic Acids Res 47, D506–D515

72. Schrodinger, LLC. (2015) The PyMOL Molecular Graphics System, Version 1.8.

73. Waterhouse, A., Bertoni, M., Bienert, S., Studer, G., Tauriello, G., Gumienny, R., Heer, F. T., de Beer, T. A. P., Rempfer, C., Bordoli, L., Lepore, R., and Schwede, T. (2018) SWISS-MODEL: homology modelling of protein structures and complexes. Nucleic Acids Res 46, W296–W303

74. Chung, C. T., Niemela, S. L., and Miller, R. H. (1989) One-step preparation of competent Escherichia coli: transformation and storage of bacterial cells in the same solution. Proc Natl Acad Sci U S A 86, 2172–2175

75. Bradford, M. M. (1976) A rapid and sensitive method for the quantitation of microgram quantities of protein utilizing the principle of protein-dye binding. Anal Biochem 72, 248–254

76. Laemmli, U. K. (1970) Cleavage of structural proteins during the assembly of the head of bacteriophage T4. Nature 227, 680–685

77. Wilkins, M. R., Gasteiger, E., Bairoch, A., Sanchez, J. C., Williams, K. L., Appel, R. D., and Hochstrasser, D. F. (1999) Protein identification and analysis tools in the ExPASy server. Methods Mol Biol 112, 531–552

78. Swinehart, D. F. (1962) The Beer-Lambert Law. Journal of Chemical Education 39, 333

